# Evaluation of copy number burden in specific epilepsy types from a genome-wide study of 18,564 subjects

**DOI:** 10.1101/651299

**Authors:** Lisa-Marie Niestroj, Daniel P. Howrigan, Eduardo Perez-Palma, Elmo Saarentaus, Peter Nürnberg, Remi Stevelink, Mark J. Daly, Aarno Palotie, Dennis Lal, Epi25 Collaborative

**Affiliations:** Cologne Center for Genomics (CCG), University of Cologne, Cologne, 50931, Germany; Broad Institute of MIT and Harvard, Cambridge, MA 02142, USA; Institute for Molecular Medicine Finland (FIMM), University of Helsinki, Helsinki, FI-00014, Finland; Center for Molecular Medicine Cologne (CMMC), University of Cologne, Cologne, 50931, Germany; Cologne Excellence Cluster on Cellular Stress Responses in Aging-Associated Diseases (CECAD), University of Cologne, Cologne, 50931, Germany; Department of Child Neurology, Brain Center Rudolf Magnus, University Medical Center Utrecht, Utrecht, Netherlands; Department of Genetics, Center for Molecular Medicine, University Medical Center Utrecht, Utrecht, Netherlands; Genomic Medicine Institute, Lerner Research Institute, Cleveland Clinic, Cleveland, OH 44106, USA; Epilepsy Center, Neurological Institute, Cleveland Clinic, Cleveland, OH 44106 USA; Stanley Center for Psychiatric Research, Broad Institute of MIT and Harvard, Cambridge, MA 02142, USA

## Abstract

Rare and large copy number variants (CNVs) around known genomic ‘hotspots’ are strongly implicated in epilepsy etiology. But it remains unclear whether the observed associations are specific to an epilepsy phenotype, and if additional risk signal can be found outside hotspots. Here, we present the largest CNV burden and first CNV breakpoint level association analysis in epilepsy to date with 11,246 European epilepsy cases and 7,318 ancestry-matched controls. We studied five epilepsy phenotypes: genetic generalized epilepsy, lesional focal epilepsy, non-acquired focal epilepsy, epileptic encephalopathy, and unclassified epilepsy. We discovered novel epilepsy-associated CNV loci and further characterized the CNV burden enrichment among phenotype-specific epilepsies. Finally, we provide evidence for deletion burden outside of known hotspot regions and show that CNVs play a significant role in the genetic architecture of lesional focal epilepsies.

## Introduction

Characterized by recurrent and unprovoked seizures, epilepsy is the third most common neurological disorder, affecting roughly 65 million people worldwide^1^. The cause of epilepsy is unknown in many patients and can be the result of a variety of insults that perturb brain function. Along with acquired causes such as trauma, infectious diseases and autoimmune diseases, genetic variants play a major role in the disease etiology^2^. To date, approximately 100 genes have been associated with epilepsy^2,3^.

The clinical representation of epilepsy is heterogeneous and subtype classification can be challenging. The epilepsies can be grouped into four major phenotypes^4^: (1) genetic generalized epilepsies (GGE), (2) focal epilepsies with non-acquired focal epilepsies (NAFE) and lesional focal epilepsies (LFE), (3) developmental and epileptic encephalopathies (DEE), and (4) unclassified epilepsies (UE). Among all epilepsy phenotypes, the DEE group has the poorest prognosis^4,5^.

In the last decade, many genetic studies have established that single nucleotide variants can confer risk or cause epilepsy^2,6^. Disease causing de novo variants have been reported in patients with DEE^7^ and seizure susceptibility variants have been identified in GGE (for a review see^8^. Focal epilepsies have been associated with germline, somatic and mosaic pathogenic variants in e.g. *PCDH19*^9^, *LGI1*, *SCN1A* and *CHRNA4* (for a review see Helbig et al., 2016^10^) and especially in genes associated with the mechanistic target of rapamycin (mTOR) pathway^11,12^.

Additionally, rare copy number variants (CNVs) are strongly implicated in the etiology of epilepsy. Around four to eight percent of DEE patients carry pathogenic CNVs^13,14^ and CNVs at genomic hotspots such as 15q13.3, 15q11.2, 16p11.2, 16p13.11 and 22q11.2 have been associated with GGE^15–22^. Rare genic CNVs were found in ~10% of GGE patients^13,18,23^ and CNVs greater than one megabase (Mb) were significantly enriched in patients compared to controls^13,14,17,24^. Deletions at 15q13.3, 15q11.2 and 16p13.11 are rarely seen in patients with DEE, highlighting the notion that the major phenotypes of epilepsy have different genetic architectures^25^. Non-recurrent deletions in *RBFOX1* have been additionally found in patients with focal epilepsies^26^ and the 16p13.11 deletion was found in a study including GGE, NAFE, and LFE patients combined^14^. However, no significant CNV association has been identified to date with NAFE^22^ and the role of CNVs in LFE has not been studied at large scale.

To date, all of the current epilepsy CNV associations have been identified through candidate loci screens, as genome-wide scans were under-powered to confirm significant genetic associations of low frequency CNVs (<1%) with epilepsy. In addition, the vast majority of CNV association studies have focused on deletions and not duplications. Lastly, no large-scale study uniformly processed or analyzed several types of epilepsy with the same genotyping platform and analysis protocol, which would enable robust comparisons across epilepsy phenotypes.

Here, we performed a large genome-wide analysis and the first CNV breakpoint association analysis of both deletions and duplications in five different epilepsy phenotypes (n=11,246 cases and 7,318 controls), to decipher epilepsy phenotype-specific patterns as well as to discover novel epilepsy-associated CNV loci.

## Methods

### Sample Ascertainment

#### Patients

Epilepsy patients and associated clinical data (n = 13,454) were ascertained from clinics distributed throughout Europe (37 sites), North America, Oceania and Asia as part of an ongoing collaborative effort by the Epi25 Consortium. Subjects were assessed for a diagnosis of developmental and epileptic encephalopathies (DEE), genetic generalized epilepsy (GGE), non-acquired focal epilepsy (NAFE), lesional focal epilepsy (LFE), with all patients having a nonspecific epilepsy diagnosis defined here as unclassified epilepsy (UE).

DEE comprised subjects with severe refractory epilepsy of unknown etiology with developmental plateauing or regression, no epileptogenic lesion on MRI, and with epileptiform features on EEG. As this is the group with the largest number of gene discoveries to date, we encouraged inclusion of those with negative epilepsy gene panel results, but we did not exclude those without prior testing.

Diagnosis of GGE required a history of generalized seizure types (generalized tonic-clonic, absence, or myoclonus seizures) and generalized epileptiform discharges on EEG. We excluded cases with evidence of focal seizures, or with moderate to severe intellectual disability and those with an epileptogenic lesion on neuroimaging (although neuroimaging was not obligatory). If EEG was not available, then only cases with an archetypal clinical history as judged by the phenotyping committee (e.g. morning myoclonus and generalized tonic-clonic seizures) were accepted.

Diagnosis of NAFE required a convincing history of focal seizure types, an EEG with focal epileptiform or normal findings, and neuroimaging showing no epileptogenic lesion or hippocampal sclerosis (MRI was preferred but CT was accepted). Exclusion criteria were a history of primarily generalized seizures or moderate to severe intellectual disability.

LFE compromised subjects with a convincing history of focal seizure types, an EEG with focal epileptiform or normal findings, and neuroimaging showing an epileptogenic lesion such as a low-grade brain tumor or a focal cortical dysplasia.

Patients with an UE diagnosis did not fulfill criteria for any of the aforementioned epilepsy phenotypes due to absence of critical data or conflicting data and are therefore under review or were labeled excluded.

Patients or their legal guardians provided signed informed consent according to local national ethical requirements. This study was approved by the institutional review boards of all participating sites (see Supplement). Samples had been collected over a 20-year period in some centers, so the consent forms reflected standards at the time of collection. Samples were only accepted if the consent did not exclude data sharing (see details in exome study using similar patient cohort https://www.biorxiv.org/content/early/2019/01/21/525683.full.pdf). Part of the dataset was published in dbGaP (phs001489.v1.p1).

#### Controls

Additional control subjects (n = 12,857) were obtained from three external large-scale genetic studies, specifically selected because genotyping was performed on the same genotyping array (Illumina Infinium Global Screening Array) and at the same center (Broad Institute) as the epilepsy cases. Controls provided as part of this study: 1) Genomic Psychiatry Cohort (GPC) controls, 2) FINRISK controls and 3) Helmsley Irritable Bowl Disease (IBD) cases and controls. For detailed description see Supplement.

### Genotyping

Samples selected for this study were all genotyped on the GSA-MD v1.0 (Illumina, San Diego, CA, USA) in separate batches. A total of 688,032 markers were used for quality control (QC).

### Genotype Sample QC

To correct for population stratification, we performed an initial round of QC based on SNP genotype data for 13,420 epilepsy cases and 12,857 controls. Samples with a call rate < 0.96 or discordant sex status were excluded. We filtered autosomal SNPs for low genotyping rate (> 0.98), case-control difference in minor-allele frequency (> 0.05), and deviation from Hardy-Weinberg equilibrium (HWE, p-value <= 0.001) before pruning SNPs for linkage disequilibrium (--indep-pairwise 200 100 0.2) using PLINK v1.9^27^ in order to perform Principal Component Analysis (PCA) to assess for population stratification. Samples with non-European ancestry were excluded based on visual clustering of the PCA.

### CNV Calling

We focused only on autosomal CNVs due to higher quality of CNV calls from nonsex chromosomes^28^. We created GC wave-adjusted LRR intensity files for all samples using PennCNV, and employed PennCNV’s CNV calling algorithms^29^ to detect CNVs in our dataset. We generated a custom population B-allele frequency file before calling CNVs. Adjacent CNV calls were merged if the number of intervening markers between them was less than 20% of the total number when both segments were combined.

### Intensity Sample QC

Intensity-based QC was conducted to remove samples with low quality data based on the following empirically defined thresholds across three different metrics: Thresholds for (1) waviness factor, (2) Log-R ratio standard deviation, and (3) B-allele frequency drift were calculated by taking the median +3x SD to determine outlying samples as performed in Huang et al.^30^. Following intensity-based QC, all samples had an Log-R ratio standard deviation of < 0.25, absolute value of waviness factor < 0.04, and a B-allele frequency drift < 0.007.

### CNV-load Sample QC

We performed a final round of sample QC by removing additional samples with excessively high CNV load based on the total number of CNV calls (>100). This threshold was determined empirically by visual inspection of distributions across all datasets combined. Our final dataset after sample QC compromised 18,564 samples: 11,246 epilepsy cases and 7,318 controls (DEE = 1,315; GGE = 3,637; LFE = 1,267; NAFE = 4,520; UE = 507).

### Call Filtering and Delineation of Rare CNVs

CNV calls were removed from the dataset if they spanned less than 20 markers, were less than 20Kb in length, had a SNP density < 0.0001 (amount of markers/length of CNV) or overlapped by more than 0.5 of their total length with regions known to generate artifacts in SNP-based detection of CNVs^31^. This included immunoglobulin domain regions, telomeric regions (defined as 500Kb from the chromosome ends), and centromeric regions (coordinates were provided by PennCNV for hg19). Further, we excluded CNVs overlapping > 80% of regions known to be recurrent copy number variations in the general population (11,732 CNVs from http://dgv.tcag.ca/dgv/app/home) for a part of the analyses (see “CNV Burden Analysis”). Additionally, all CNV calls spanning more than 20 markers and equal to or more than 1Mb in length were included in the analysis even if the SNP density was < 0.0001^30,31^.

We assigned all CNV calls a specific frequency count using PLINK v.1.07^32^, with the option -- cnv-freq-method2 0.5. Here, the frequency count of an individual CNV is determined as 1 + the total number of CNVs overlap by at least 50% of its total length (in bp), irrespective of CNV type. We then filtered our callset for rare CNVs with MAF < 1% (a frequency of 186 or lower across 18,564 samples).

After CNV quality control, 12,765 of 18,564 (7,748 cases and 5,017 controls) QC-passed individuals had one or more rare CNVs.

### CNV Annotation

CNVs were annotated for gene content and recurrent deletion hotspots for epilepsy and neurodevelopmental disorders (NDD) with various annotation files including gene name and the corresponding coordinates in hg19 assembly using in-house perl scripts (available on request). We annotated 89 genes that were previously associated with epilepsy^2,3^, 93 genes associated with NDD^33^, 2,680 genes intolerant for protein truncating variants defined as pLI > 0.95^34^ (probability of loss-of-function intolerance [pLI] score > 0.95), >28,000 annotated regions from UCSC refseq genes, eight recurrent hotspot deletion regions for epilepsy and six recurrent hotspot regions for NDD^35^. We only considered a CNV as “coding” if it overlapped 80% of a gene^36^. We considered all other CNVs as “non-genic”.

Cytogenic testing is well-established for diagnostic evaluation of patients with neurodevelopmental disorders including epilepsies. It is generally established that large deletions, deletions intersecting haploinsufficient genes, and large duplications are considered as likely pathogenic for epilepsy^37^. Therefore, we considered a CNV as “likely pathogenic” as defined by ACMG guidelines^38^, i.e. if its length exceeded 2Mb, it overlapped a known hotspot region for epilepsy, a gene with pLI > 0.95, or a known epilepsy-associated gene.

### CNV Burden Analysis

We measured CNV burden for all five epilepsy phenotypes using three separate categories to evaluate relative contribution on epilepsy type risk: (1) the total length of all rare CNVs within an individual (CNV length), (2) the carrier status of rare CNVs intersecting genes and neurodevelopmental or epilepsy associated CNVs hotspot regions, and (3) the carrier status of rare likely pathogenic CNVs. For length and CNV burden in different gene and hotspot lists, deletions and duplications were analyzed separately. For likely pathogenic CNV burden duplications and deletions were analyzed according to the definition of “likely pathogenic” CNVs mentioned before. To assess for a CNV burden difference between epilepsy cases and controls, we fitted a logistic binomial (for hotspot regions including CNVs from the general population) or Poisson (for gene lists and likely pathogenic CNV burden excluding CNVs from the general population) regression model using the “glm” function of the stats package (https://github.com/SurajGupta/r-source/tree/master/src/library/stats/R) in R for common and rare CNVs respectively^30^:

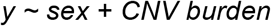

where ‘y’ is a dichotomous outcome variable (epilepsy type = 1, control = 0); ‘sex’ is used as a covariate and ‘CNV burden’ represents one of the categories mentioned above. For all burden analyses, ORs, 95% confidence intervals (CIs), and significance were calculated. ORs were calculated by taking the exponential of the logistic regression coefficient. ORs above one indicate an increased risk for the specific epilepsy type per unit of CNV burden. Significance threshold was corrected for multiple testing using Bonferroni correction. Bonferroni multiple-testing threshold for significance was calculated combined for all epilepsy phenotypes and CNV types for all three categories ((1) CNV length burden p < 1.6e-3; (2) genome-wide burden p < 8.33e-4; (3) likely pathogenic CNV burden p < 0.01).

### Regression of Potential Confounds on Case-Control Ascertainment

It is important to ensure that any bias in gender and ancestry does not drive spurious associations with epilepsy. To ensure the robustness of the analysis, CNV burden analyses included potential confounding variables as covariates in a logistic regression framework. Due to the number of tests run at breakpoint level association, we employed a step-wise logistic regression approach to allow for the inclusion of covariates in our case-control association, as previously described in Marshall and Howrigan et al.^31^, which we term the epilepsy residual phenotype. Covariates included sex for burden and breakpoint association analysis and the first ten ancestry principal components for breakpoint association analysis.

To calculate the epilepsy residual phenotype, we first fitted a logistic regression model of covariates to affection status, and then extracted the Pearson residual values for use in a quantitative association design for downstream analyses. Residual phenotype values in cases are all above zero, and controls below zero, and are plotted against overall Kb burden in Figure S1.

### CNV Breakpoint Level Association

The CNV breakpoint level association was performed by quantifying the frequencies of case and control CNV carriers at all unique CNV breakpoint locations (i.e., the SNP probe defining the start and end of the CNV segment); the full set of CNV breakpoints represents the genome-wide space of CNV variation between cases and controls.

CNV breakpoint level association was run using the epilepsy residual phenotype as a quantitative variable, with significance determined through 1,000,000 permutations of phenotype residual labels using PLINK v1.07^32^. An additional z-scoring correction was used to efficiently estimate two-sided empirical *p*-values for highly significant loci. A fraction of our controls were patients from an Irritable Bowl Disease (IBD) project, and therefore to rule out confounding, we ran the same CNV breakpoint level association for the “IBD-controls” from the Helmsley dataset (since these represent IBD cases) and used them as cases to test association using the remaining controls as comparison group. IBD-related CNV breakpoints with *p*-values <10e-3 after genome wide correction were removed from the combined analysis (epilepsy cases vs all controls including IBD fraction). Association tests were conducted for all CNV types, deletions, and duplications independently. CNVs spanning the centromere were merged to one. Bonferroni correction for multiple testing was used to identify significance threshold. Loci that surpassed genome-wide multiple testing correction in either test were followed up by manual CNV quality evaluation: B-allele frequency and LogR-ratio were manually investigated using perl scripts provided by PennCNV and UCSC genome browser hg19 (https://genome.ucsc.edu/).

### Phenotype Analysis

The phenome-wide association study (PheWAS) design requires a good signal to noise ratio to discover novel CNV associations. To enrich for high confidence pathogenic CNVs, we tested the burden of big CNVs (>2Mb) in patients with a specific phenotype among the different epilepsy phenotypes. Based on the data collected through the Epi25 consortium, we were able to include 43 different phenotype categories in the PheWAS (see Supplementary Methods). *P*-values and ORs were obtained using a Fisher’s Exact Test (two-sided). Multiple testing correction for 161 tests results in a significant p-value < 3.1*10^-4^. We performed a meta-analysis for the association of GGE patients with big duplications (> 2 Mb) with febrile seizures to exclude a possible center bias using the R package “metafor” (https://cran.r-project.org/web/packages/metafor/metafor.pdf).

## Results

### Elevated epilepsy type-specific CNV burden in DEE and GGE patients

We applied logistic regression to investigate whether the five epilepsy phenotypes have on average a greater genomic region covered (combined CNV length) by either deletions or duplications. After correction for 30 tests, we found that patients with DEE and GGE showed independent enrichment for total deletions of an overall length of >2Mb compared to controls (DEE: OR 2.91 [1.63-4.72], p = 7.13e-5; GGE: OR 1.85 [1.27-2.58], p = 6.5e-4) (Figure 1A). UE was the only epilepsy type with significant burden for duplications of an overall length of >2Mb (OR 3.85 [2.71-5.3], p = 2.63e-15; Figure 1B).

**Figure 1.**
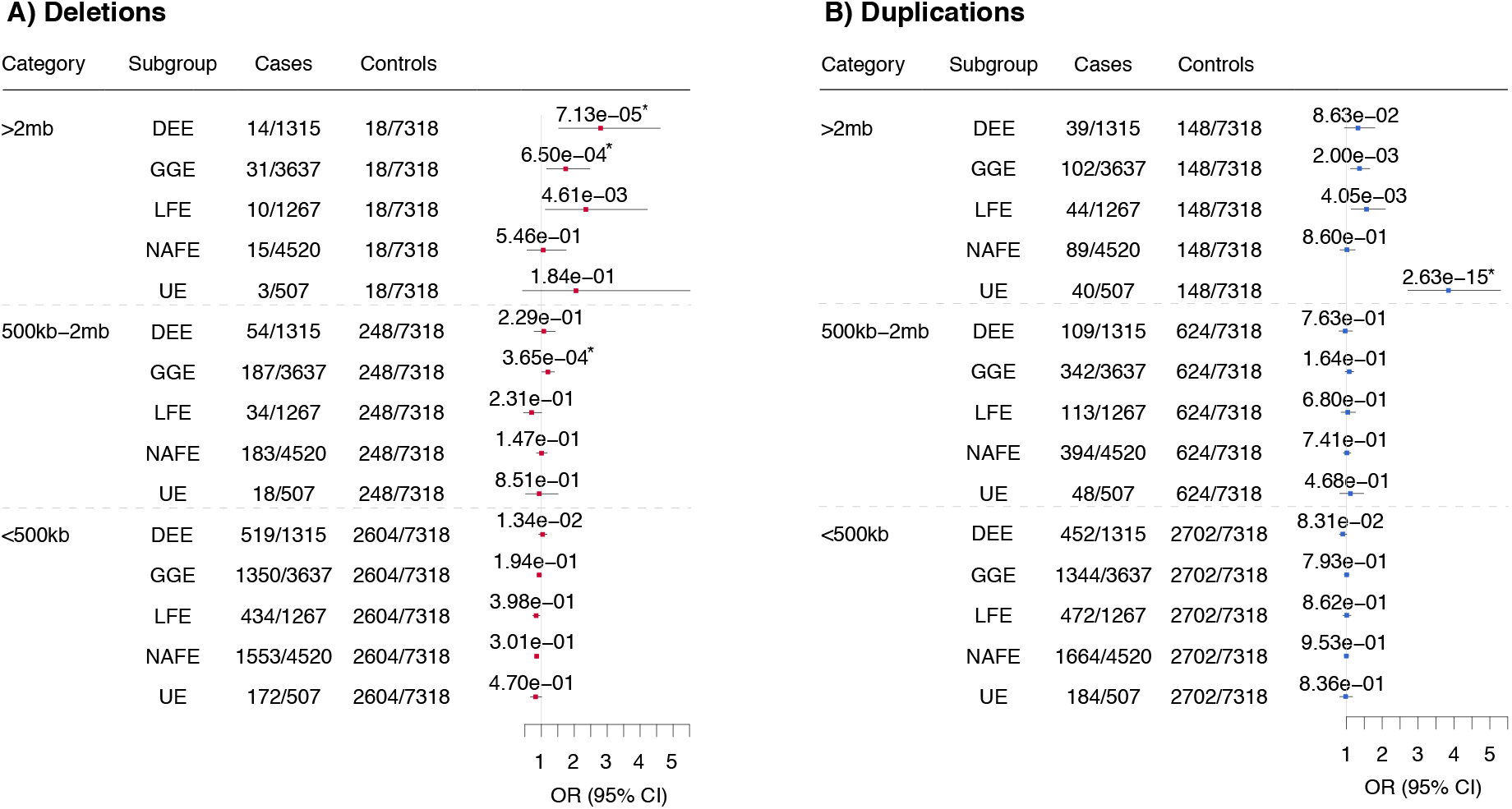
Global burden of CNV by overall length across five epilepsy types. Rare CNV burden observed in the different epilepsy types is shown for (A) deletions and (B) duplications. Odds ratios (ORs) and p-values were calculated using a Poisson logistic regression for rare CNVs with sex as a covariate in three different categories (overall genomic sequence loss in one individual of >2Mb, 500Kb-2Mb and <500Kb). DEE = Developmental and epileptic encephalopathies; GGE = Genetic generalized epilepsies; LFE = Lesional focal epilepsies; NAFE = Non-acquired focal epilepsies; UE = Unclassified epilepsies; * = p values surpassing the Bonferroni multiple testing for 30 tests cut-off (*p< 1 .63^-3^).

### Enrichment of gene-sets and CNV hotspots in DEE, GGE, and NAFE patients

Next, we measured if the CNV burden was concentrated within defined sets of genes and known deletion hotspots for epilepsy (Epi) and neurodevelopmental disorders (NDD). Compared to deletions identified in the controls, we found that the epilepsy hotspot list, genes intolerant for truncating variants, and coding regions were enriched for patient deletions (Figure 2). DEE and GGE patients showed a significant burden of deletions in genes with pLI > 0.95 (DEE: OR 1.85 [1.3-2.53], p = 2.78e-4; GGE: OR 1.58 [1.28-1.91], p = 7.2e-6). Additionally, GGE patients showed an enrichment of deletions at previously identified epilepsy hotspots (OR 5.21 [3.59-7.7], p = 2.01e-17) and in coding regions (OR 1.15 [1.07-1.24], p = 2.35e-4) but no significant enrichment of known epilepsy genes. Furthermore, we detected a significant deletion enrichment in NAFE patients at previously identified epilepsy deletion hotspots (OR 2.42 [1.61-3.69], p = 2.87e-5). In contrast, no enrichment was observed in any genes or loci tested when duplications were considered in any epilepsy phenotype (Figure S2).

**Figure 2.**
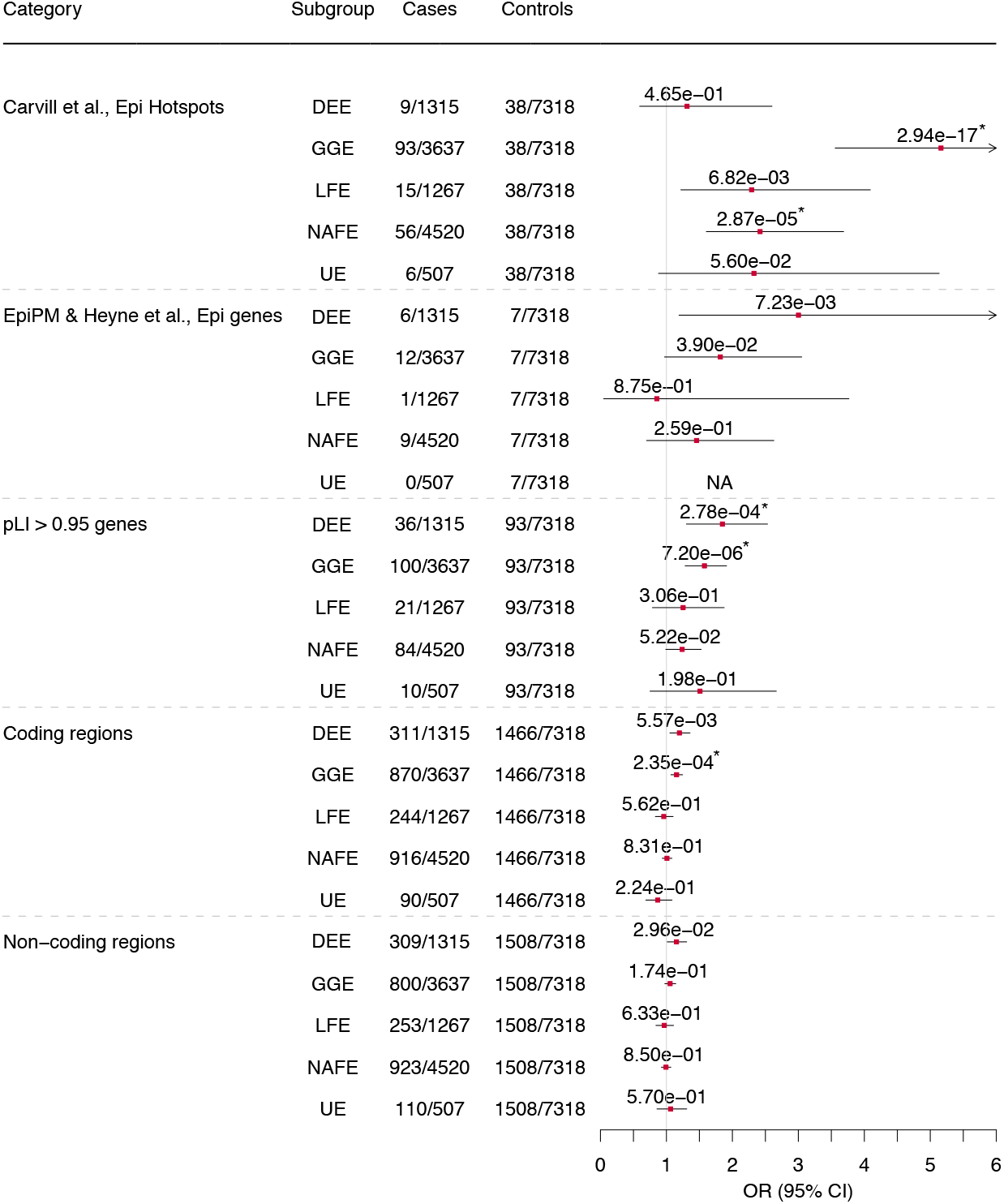
The global burden of deletions across different gene sets, hotspot regions and non-coding regions in five different epilepsy phenotypes. Common deletion burden was elucidated for epilepsy hotspot regions^35^ and rare (< 1% frequency) deletion burden was elucidated for all other gene lists (Category). Odds ratios (ORs) and p-values were calculated using a binomial regression for common CNVs and a Poisson regression for rare CNVs with sex as a covariate. CNVs are defined as “genic” if they overlap 80% of a gene. Notably, not all individuals carry a CNV. (Results of CNV burden in NDD hotspots and NDD genes are not shown due to very small sample sizes and no significance; results of duplication burden are shown in Supplementary Figure 2). 95% CIs are clipped to arrows when they exceed a specified limit. DEE = Developmental and epileptic encephalopathies; GGE = Genetic generalized epilepsies; LFE = Lesional focal epilepsies; NAFE = Non-acquired focal epilepsies; UE = Unclassified epilepsies; * = p values surpassing the Bonferroni multiple testing for 60 tests cut-off (*p < 8.33e^-4^).

### Enrichment of likely pathogenic CNVs in all epilepsy phenotypes

For our next category, we evaluated the combined burden of the CNVs that are considered in the literature as ‘likely pathogenic’ (according to ACMG, see “Methods” for selection criteria) in the five studied epilepsy phenotypes. Likely pathogenic CNVs were identified in 6.08 % of DEEs, 7.67 % of GGEs, 5.92 % of LFEs, 4.67 % of NAFEs, and 9.27 % of UEs. However, likely pathogenic CNVs were also present in 3.56 % of controls. Nevertheless, in a direct comparison with the controls, we observed a significant enrichment of likely pathogenic CNVs in all epilepsy phenotypes (Figure 3). The likely pathogenic CNV effect size was greatest in patients with UE (OR 2.63 [1.92-3.52], p = 4.16e-10; Figure 3), mainly driven by large duplications (Figure 1B).

**Figure 3.**
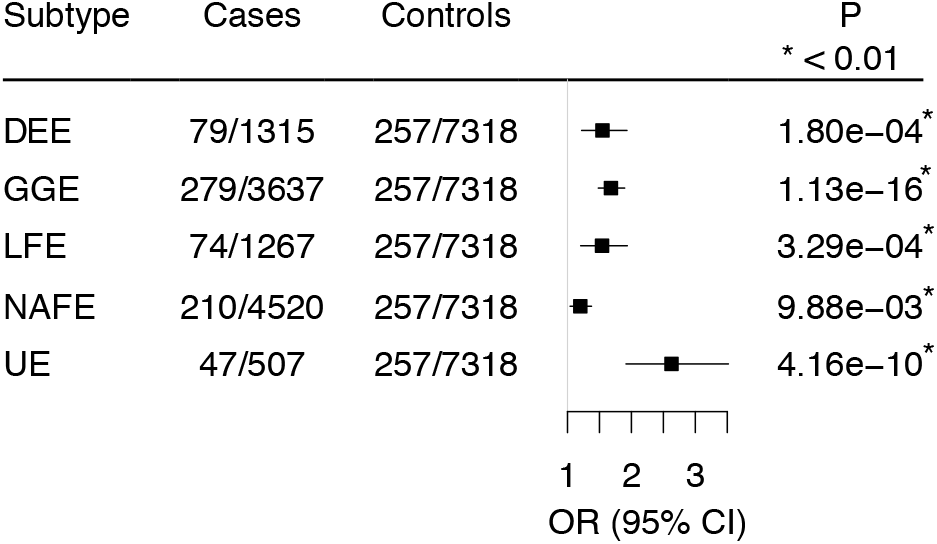
Global burden of likely pathogenic CNVs across five different epilepsy phenotypes. Likely pathogenic CNVs were defined as frequency < 1%, >=2Mb, deletions in known epilepsy hotspots, deletions in known epilepsy genes, or deletions in genes with pLI > 0.95. Odds ratios (ORs) and p-values were calculated using a poisson logistic regression for rare CNVs with sex as a co-variable. Genic CNVs are defined as those that overlap 80% of any exon of a known protein-coding gene. DEE = Developmental and epileptic encephalopathies; GGE = Genetic generalized epilepsies; LFE = Lesional focal epilepsies; NAFE = Non-acquired focal epilepsies; UE = Unclassified epilepsies; * = p values surpassing the Bonferroni multiple testing for five tests cut-off (*p < 0.01).

### Genome-wide CNV breakpoint association reveals significant loci outside of known hotspot regions

In total, five independent CNV loci in five epilepsy phenotypes surpassed genome-wide significance; four loci have been previously reported in association with GGE^15–18^ and one has never been associated with epilepsy before. For three of the identified CNV loci we extended the phenotypic spectrum by identifying novel epilepsy phenotype associations. In line with previous results from candidate loci studies, our analysis showed that patients with GGE were most significantly enriched for deletions overlapping hotspot loci on chromosomes 15q13.2-q13.3 (p = 2.55e-08) and 16p13.11 (p = 3.43e-08; Figure 4A, Figure S4). We identified a duplication association with GGE that was located on chromosome 9, spanning 9p11.2, the centromere and 9q21.11 (p = 1.53e-07; Figure 4B, Figure S4, S5), a locus associated for the first time with an epilepsy phenotype. The DEE analysis revealed a genome-wide significant duplication locus overlapping the recurrent region on chromosome 15q11.2-q13.1 also known as the Prader-Willi/Angelman critical region (p = 2.15e-10; Figure 4B). No locus was significantly enriched in the NAFE cohort. Deletions in LFE patients were enriched at epilepsy hotspot 16p13.11 (p = 7.08e-08; Figure 4A), and duplications also at 9p11.2-9q21.11 (p = 1.09e-10; Figure 4B; Figure S4, S5). Finally, the UE association analyses identified significant enrichment for duplications at 1q21.1 and 9p11.2-9q21.11 (p = 3.30e-11; p = 3.37e-18; Figure 4B). To verify the novel duplication region 9p11.2-9q21.11 significantly enriched in GGE, LFE and UE patients, we plotted the Log-R Ratio (LRR) intensity and B-Allele Frequency (BAF) of the probe-levels for a subset of six patients in Figure S5.

**Figure 4.**
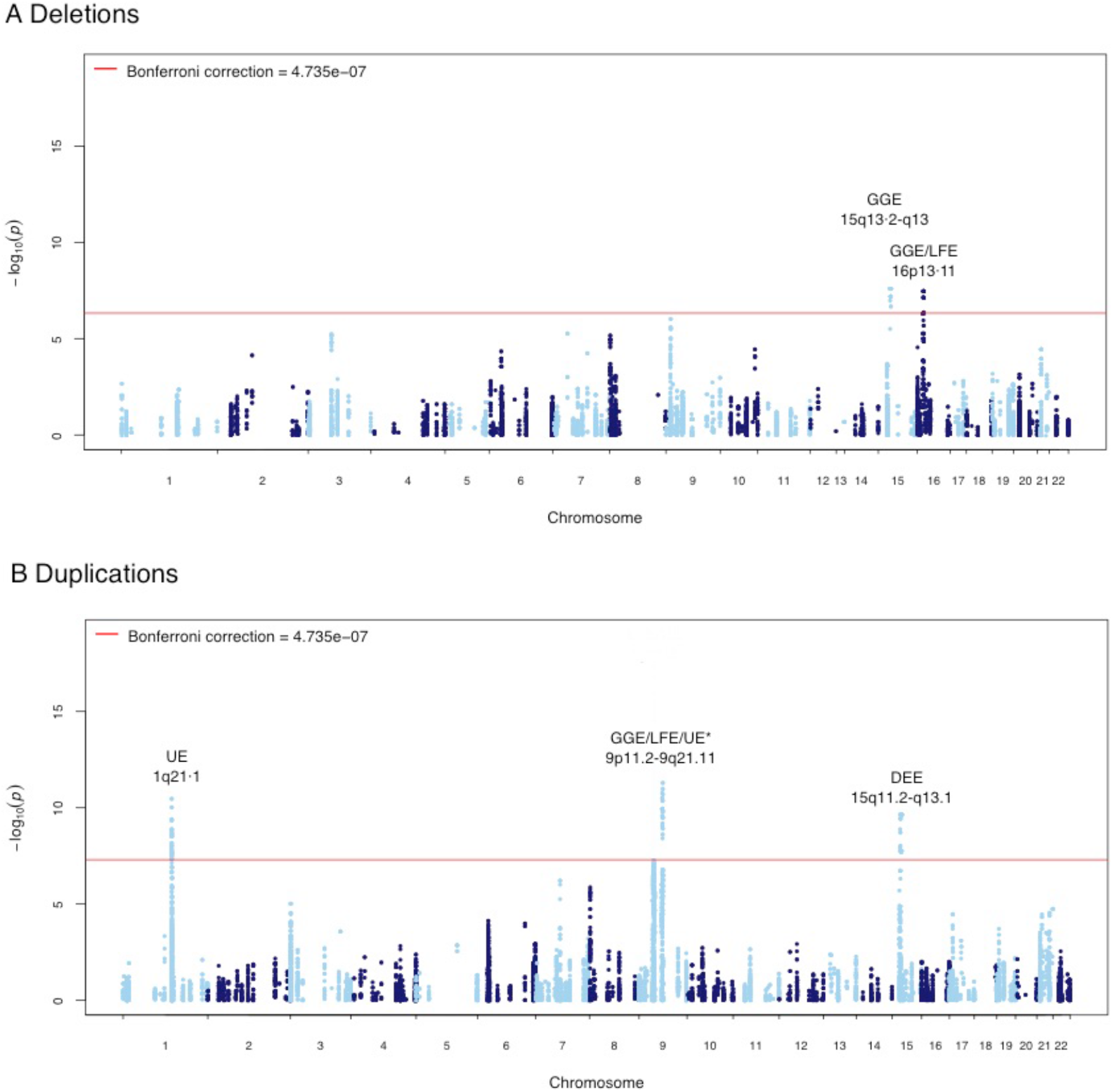
Genome-wide CNV breakpoint association. Manhattan plot displaying the –log10 deviance p-value for A) Genome-wide deletion breakpoint association for DEE, GGE, LFE, NAFE, and UE and B) Genome-wide duplication breakpoint association for DEE, GGE, LFE, NAFE, and UE. P-value cutoffs corresponding to correction for 105,596 tests at 4.743e-7 are highlighted in red. Loci significant after multiple test correction in the appropriate epilepsy type are labeled. * = Two association signals of the same duplication identified by the start- and end-breakpoint at 9p11.2 and 9q21.11 in GGE, LFE, and UE. DEE = Developmental and epileptic encephalopathies; GGE = Genetic generalized epilepsies; LFE = Lesional focal epilepsies; NAFE = Non-acquired focal epilepsies; UE = Unclassified epilepsies.

### PheWAS analysis reveals enrichment of large CNVs (> 2Mb) in epilepsy subtypes

We performed a phenome-wide association study (PheWAS) to identify an association between large effect CNVs and a large number of different phenotypes. We analyzed whether the CNV burden is enriched in any clinical phenotype within the five different epilepsy phenotypes. After multiple testing correction for 161 applied tests, we identified two significant associations. We observed a 3.25-fold enrichment of large duplications (> 2Mb) in patients with GGE and febrile seizures when comparing to GGE patients without febrile seizures (OR 3.25 [1.8-5.92], p = 4.07e-05; Table S2). Further, a 2.72-fold enrichment of large duplications was detected for focal epilepsy patients with structural abnormalities versus without (OR 2.72 [1.57-4.56], p = 2.33e-04; Table S2). An evaluation of types of lesions in this group showed that pathogenic CNVs are not specific to a single lesion type but found in patients with five different lesion types (Figure S6).

## Discussion

In this study, we identify several novel CNV-epilepsy associations using a case-control approach with >18,000 individuals genotyped on the same platform and analyzed with the same CNV calling, quality control, and analysis pipeline. We observe an increased burden of CNVs in different epilepsy phenotypes, report novel risk loci that surpass genome-wide multiple testing correction, and show that also LFE can be associated with an increased CNV burden. Consistent with results from genetic studies in other neurodevelopmental disorders, we show that novel risk loci lay at the ultra-rare end of the CNV frequency spectrum. Thus, larger samples will be needed to identify additional risk loci at convincing levels of statistical evidence^30,31^.

### CNV Burden

We and others have previously shown a burden of deletions overlapping genes associated with neurodevelopmental processes in patients with GGE, and that the signal was particularly concentrated within epilepsy hotspot loci^15–22^. In the present study we were able to replicate the original GGE signal with a significant enrichment for deletions in epilepsy hotspots. Additionally, we observed a significant deletion burden in genes intolerant for protein truncating variants in the general population, which has been suggested recently in a smaller cohort of 160 generalized, 32 focal, and six unclassified epilepsy patients^39^. Consistent with the well-established role of rare, large effect CNVs in the etiology of the severe and early onset DEEs^13^, we identified a significant deletion enrichment covering genes intolerant for truncating variants in the general population. Previous studies did not find significant differences between focal epilepsy patients and controls within hotspot loci, most likely due to the small sample size^22^. Here, we detect deletions overlapping epilepsy hotspot regions enriched in patients with NAFE. We observed enrichment for overall large duplications burden (>2Mb) for 6% of patients with UE, although we cannot exclude that a subset of patients may have a severe neurodevelopmental disease phenotype. This proportion is lower than in previous reports that identified that 15-20% of individuals with unexplained neurodevelopmental disorders carry pathogenic CNVs^40^. Although epilepsy associated brain lesions have mainly been associated with somatic variants, which affect the mechanistic target of rapamycin (mTOR) pathway^11,12^ also germline variants in *DEPDC5* have been identified as risk factors for lesional epilepsies. Here, we show that CNVs play a role in the etiology of LFE. The detected pathogenic CNVs were not specific to a single brain lesion, suggesting that the CNVs confer risk to the epilepsy rather than to the lesion itself.

CNVs are present in most people and usually represent benign genetic variation without clinical significance^41^. Therefore, we concentrated on the burden of likely pathogenic CNVs that were 1.2-2.61-fold enriched in epilepsy patients. Although we used state-of-the-art criteria to support the categorization as ‘likely pathogenic’ CNV, the modest enrichment indicates that many population controls carry similar types of CNVs. This observation is in accordance with the presence of recurrent CNVs in epilepsy hotspot loci in healthy controls, suggesting an incomplete penetrance for epilepsy risk (Dibbens et al., 2009, Crawford et al., 2018). Additionally, detection of large gene-disrupting CNVs and epilepsy-associated gene deletions does not imply causality but rather increased susceptibility or incomplete penetrance. Many CNV hotspots and large-gene disrupting CNVs are known to be co-morbid with other disorders like intellectual disability (Mullen et al., 2013) and autism^42–45^, but we did not observe an enrichment of likely pathogenic CNVs in patients with these comorbidities in our cohort (data not shown). Interestingly, we found an enrichment of large duplications (>2Mb) in GGE patients with febrile seizures compared to GGE patients without febrile seizures (Table S2, Figure S3). Additional comorbidities in GGE patients with CNVs have been reported before (Mullen et al., 2013). Large duplications at 1q21.1, 22q11.2, and 16p11.2 are known to be enriched in syndromic epilepsies^46–48^, suggesting that those GGE patients carry additional phenotypic co-morbidities.

### Genome-wide CNV breakpoint association

Several recurrent CNVs have been previously associated with epilepsy^15,16^, however all have been identified in candidate loci studies. In this study, our sample size and uniform CNV calling pipeline allowed us to test CNV loci at genome-wide scale with adequate power at the CNV breakpoint level. Here, we performed the first genome-wide CNV breakpoint association analysis to identify associated loci among different epilepsy phenotypes. We replicated four of seven previously published locus-associations with epilepsy types at genome-wide significance level (1q21.1, 15q11.2, 15q13.3 and 16p13.11)^15–18^, whereas 16p11.2, 16p12, and 22q11.2 only reached suggestive significance (p-value < 0.05), suggesting that larger datasets are needed to reach genome-wide significance. The majority of these previously established loci are co-morbid with other neurodevelopmental disorders such as schizophrenia, psychotic disorder, autism or intellectual disability^31,49,50^. Notably, our previous GGE CNV study re-evaluated clinical records of GGE patients carrying a 22q11.2 deletion, revealing additional congenital and developmental features^17^. Possibly in this study, we used more stringent sample inclusion criteria with a smaller fraction of patients with comorbidities. This may explain why three out of seven recurrent loci were not significantly enriched in our analysis. Nonetheless, we show a significant association of deletions in 16p13.11 with LFE. Previously, deletions of 16p13.11 were found to be enriched in candidate loci studies of GGE and CECTS (Childhood epilepsy with centrotemporal spikes) along with autism, intellectual disability, schizophrenia and additionally in non-lesional focal epilepsies^15,18^. The signal of non-lesional focal epilepsies could have been driven by misdiagnosed patients with small lesions undetectable by neuroimaging so that a lesional focal epilepsy might not have been confidently ruled out in these patients.

GGE, LFE and UE were associated with a genome-wide significant duplication spanning 9p11.2, the centromere and 9q21.11, which has never been associated with epilepsy before. Both loci harbor genes highly expressed in the brain (9p11.2: FAM27E3; 9q21.11: e.g. PIP5K1B, APBA1). However, regions around the centromere of chromosome 9 (9p12, 9q13-q21.12) have also been repeatedly found and described as euchromatic cytogenetically visible copy number variations (CG-CNVs)^51,52^ in close proximity to the regions we identified. So far, these regions have been reported to be prone to benign CNVs and have not been associated with any phenotypic consequence before. Further large-scale studies will help to confirm this signal (see also Figure S5 for examples of CNVs at this region). CNVs covering the identified region and additional genomic regions have been associated with several severe syndromes. Among patients with 9p duplication syndrome characterized by growth and developmental delay^53^, a patient duplication covering 9p11.2 was described^54^. Typical characteristics for the 9p duplication syndrome include further microbrachcephaly, atypical face morphology, and delayed bone age^55–57^. Wilson and colleagues proposed that the spectrum of clinical severity in the 9p duplication syndrome roughly correlates with the extent of trisomic chromosome material (Wilson et al., 1985), which could explain a milder phenotype for our LFE and UE patients with duplication of loci 9p11.2 and not the entire chromosome arm. The 9p11.2-9q21.11 duplication is enriched in epilepsy patients similar to the 15q.11.2 deletion, as it is present in the general population but clearly enriched in people with various neuropsychiatric disorders and idiopathic generalized epilepsies implicating that this CNV acts as a risk factor instead of a large effect variant.

### Study limitations

It is important to note that CNV breakpoints in the current study are estimated from genotyped SNPs around the true breakpoint, and these breakpoint estimates are limited by the resolution of the genotyping platform. Last, we recognize that especially small structural variants are not detectable with current genotyping platforms^58^. New technologies for whole-genome sequencing will ultimately enable the assessment of the contribution of a wider array of rare variants, including balanced re-arrangements, small CNVs^59^ and short tandem repeats^60^.

### Summary

Large-scale collaborations in epilepsy genetics have greatly advanced discovery through genome-wide association studies. Here, we have extended this framework to rare CNVs in five different epilepsy phenotypes including stringent ancestry and data quality control criteria, after generating the data under the same genotype array and calling pipeline for each subject. Our results help to refine the list of promising candidate CNVs associated with specific epilepsy types and extend the phenotypic spectrum for identified loci. We are confident that the application of this framework to even larger datasets has the potential to advance the discovery of loci and identification of the relevant genes and functional elements.

## Author Contribution

Conceptualization: L.M.N., D.L., E.C.;

Methodology: L.M.N., D.L., E.C.;

Software: L.M.N., E.P.P., E.S.;

Formal Analysis: L.M.N.;

Investigation: D.L., E.C.;

Resources: E.C.;

Writing – Original Draft: L.M.N., D.L.;

Writing – Review & Editing: L.M.N., R.S., P.N., E.C., D.L.;

Funding Acquisition: P.N., E.C.;

Supervision: D.L.

